# Attention-dependent physiological correlates in sleep deprived healthy humans

**DOI:** 10.1101/2020.12.05.412890

**Authors:** Valentina Cesari, Elena Marinari, Marco Laurino, Angelo Gemignani, Danilo Menicucci

## Abstract

Distinct cognitive functions are based on specific brain networks, but they are also affected by workload. The workload is a common factor affecting cognitive functioning that, by activating the Central Autonomic Network, modulates heart rate peripheral correlates of cognitive functioning. Based on these premises, we expected that the peripheral patterns associated with different attentional systems would have common (workload-related) and specific (task-dependent) components. To disentangling the components, a profile of peripheral physiological correlates of cognitive functioning was derived by studying healthy volunteers while performing different cognitive tasks during baseline and post-sleep deprivation conditions. Post-sleep deprivation condition was introduced to increase workload during tasks, allowing the investigation of the same participant at different levels of workload in different conditions. We performed, in each condition, physiological recordings of heart pulse, facial temperature and head movements during tasks assessing attentional networks efficiency (ANT - Attentional Network Task; CCT - Continuous Compensatory Tracker). We assessed perceived workload after the execution of these tasks. Physiological correlates of cognitive performance were identified by associating changes of task indices with the corresponding changes in physiological measures from baseline to post-sleep deprivation condition. Correlation analyses were performed after correction for the between-conditions workload changes: indeed, mental and physical demands of perceived workload increased after sleep deprivation. We found that alerting/vigilance has specific physiological correlates as indicated by the negative correlation between changes in ANT-alerting score and changes in amplitude of head movements and the positive one between changes in CCT-visuomotor speed indexing alertness and changes in facial temperature.

## 1. Introduction

The daily living of human beings is driven by cognitive processes related to the transformation, reduction, elaboration, storage and recovery of sensory input in the real world [1]. The functioning of distinct cognitive functions is based on specific brain networks and gives rise to distinguishable activations; however, a common factor affecting cognitive functions is workload, the multidimensional construct quantifying the level of mental and physical effort put forth by a performer in response to cognitive tasks. The evaluation of workload spans from classical neurocognitive tests to dynamic situations such as aviation and driving [2]; however, it is usually considered a property of an individual attitude toward a demanding situation rather than a task [3,4].

Workload is sustained by arousal and it is described as an indicator of pressure on working memory [5,6]. The arousal during workload implies an autonomic activation involved in non-consciously coordinated bodily responses for homeostasis) [7]. Autonomic arousal is sustained by the activity of the Central Autonomic Network [8] that includes: insula, amygdala, hypothalamus, periaqueductal gray, parabrachial complex, nucleus of the solitary tract, and the ventrolateral portions of the medulla.

Besides workload-related effects, several studies have highlighted specific neurofunctional patterns associated with cognitive tasks involving different domains. Concerning the three networks of attention, alerting has been associated with various frontal and parietal regions with strong thalamic involvement; orienting has been associated with parietal sites and frontal eye field; executive to anterior cingulate cortex and dorsolateral prefrontal cortex [9,10]. Both the areas of arousal and those of individual cognitive functions give rise to changes in peripheral autonomic outputs [11,12] that we would investigate in the present work.

From a methodological perspective, cognitive functioning has been studied by taking advantage of intra-subject variability, which has been widely assessed in the context of some stress-related states, such as sleep deprivation, that acts as a reliable paradigm of acute stress [13]. Indeed, acute and chronic sleep deprivation represents a stressful condition of modern society, posing high and significant risks for quality of life and psycho-physical wellbeing, including cognitive performance degradation [14]. Both acute total sleep deprivation and chronic sleep restriction increase homeostatic sleep - process S - leading to sleep debt. Process S augments during wakefulness and decreases during sleep time [15]: this increase impairs cognitive functions during wakefulness such as attention, cognitive speed, and memory [16]. For this reason, several studies used an acute sleep deprivation model to understand its impact on various cognitive domains [13]. Indeed, acute sleep deprivation might negatively influence some aspects of cognitive functioning, in particular vigilance, which, if lowered, increases the risk of accidents [17]. In these studies, the autonomic outputs are often assessed, since higher mental workload is associated with a decrease of parasympathetic (“rest or digest”) autonomous nervous system activity and an increase in sympathetic (“fight or flight”) activity [18, 19]. These changes in autonomous nervous system activity have been estimated with several peripheral physiological measures such as heart rate, skin conductance, and peripheral temperature [20] during tasks assessing different cognitive domains. For example, there is a positive association between cognitive load, level of glucose and oxygen in the brain [21], and forehead temperature [22, 23]. An increase in heart rate with increasing difficulty in decision making and level of attention [24–26] and as a decrease in heart rate variability with increasing difficulty in response inhibition and memory [27–29] have also been documented. Regarding skin conductance, its increase has been detected during the performance on attention, memory, vigilance, and visual tracking tasks [30–36].

In summary, many studies have used specific cognitive tasks to establish cognitive-related physiological outcomes. Several studies, taking into account subjective workload, have shown that physiological measures depend on the degree of subjectively perceived difficulty while performing tasks. This evidence implies the urge to identify the modulation of specific cognitive functions on the peripheral physiological signals, as suggested by the specific central networks sustaining the different cognitive domains. To this aim, we studied the peripheral physiological correlates in subjects undergoing different cognitive tasks at baseline and post-sleep deprivation, and we assess the perceived workload in performing the tasks in these two different conditions. While performing different tasks in the same subjects, we had the opportunity to separate the common peripheral response over the tasks, putatively related to cognitive workload, from that characterizing specific cognitive functions engaged in each specific task.

## 2. Material and Methods

### 2.1 Participants

Fifteen healthy young volunteers (9 females and 6 males; mean age ± SD: 24.5 ± 2 yrs) were enrolled for this experimental protocol. Subjects eligible for inclusion met the following criteria: absence of psychiatric symptoms as assessed by Symptom Checklist-90-Revised [37–39]; absence of sleep-wakefulness disorders as assessed by Insomnia Severity Index (ISI) [40,41] and Epworth Sleepiness Scale (ESS) [42,43]; absence of organic pathologies and psychotropic addiction as assessed by a qualitative anamnestic questionnaire.

### 2.2 Experimental protocol

The experimental protocol consisted of two sessions (Figure 1a), which were randomized and balanced across participants and which took place with a one-week interval, at least. Each session started at 6 pm and each volunteer was tested individually. Laboratory room was temperature-controlled (22 C°).

**Figure 1:**
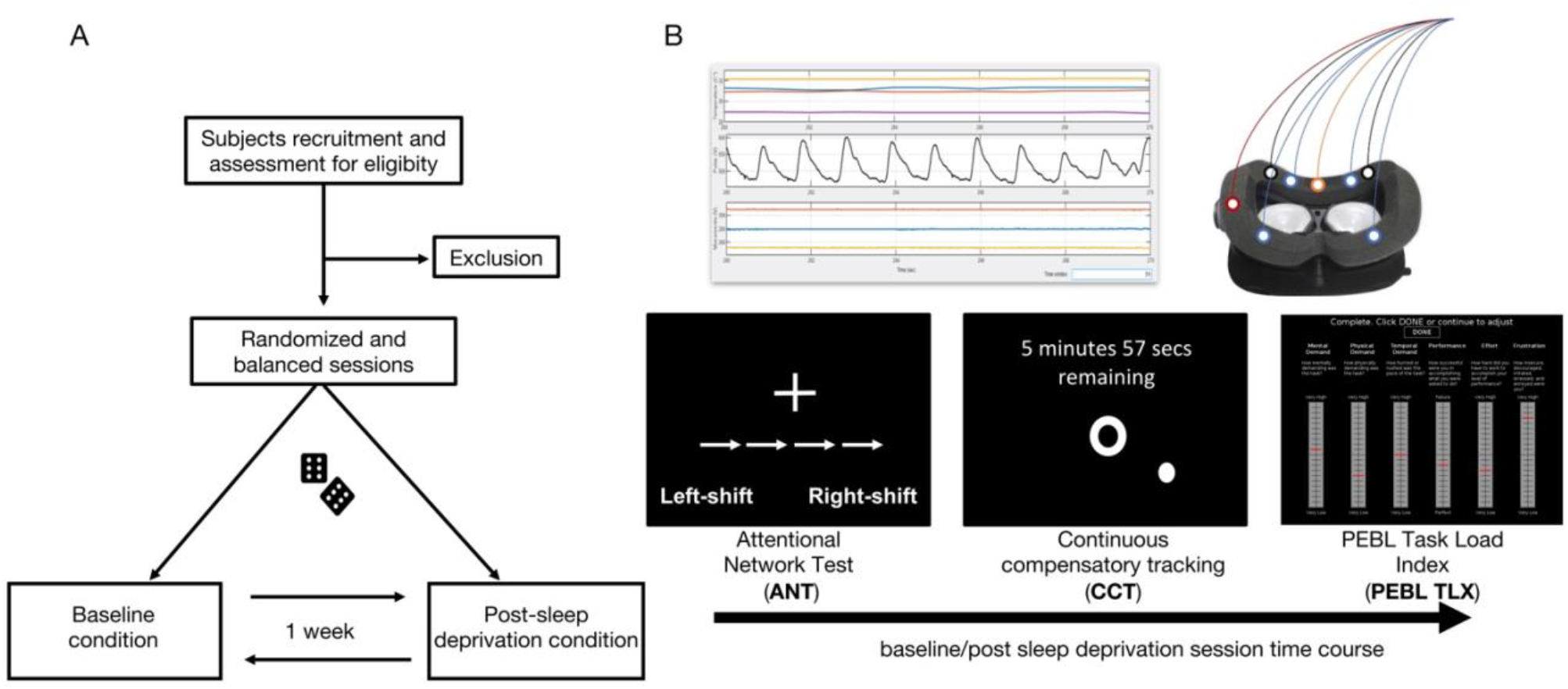
Experimental Protocol. A. Flow chart of experimental protocol. B. Above, a sketch of the online recording of facial temperature, head movements and heart pulse by means of the PERFORM system while the subject is sitting in front of PC; below, the schematic representation of the experimental timeline for both baseline and post sleep deprivation conditions.

Two cognitive tests followed by a perceived workload assessment were completed both in the baseline session and post-sleep deprivation session (Figure 1b). To ensure that volunteers had normal sleep before baseline session, sleep monitoring was accomplished by actigraphy registration and by filling out a sleep diary [44,45]. Actigraphic monitoring was also used to ensure volunteers were completely sleep deprived in post-sleep deprivation sessions; any sleep episode implied the exclusion from the study. For this double purpose, participants wore an ActiGraph wGT3X-BT (ActiGraph, Pensacola, FL, USA) placed on their non-dominant wrist. Data was analyzed and visually inspected using Actilife software (version 6.11.9).

For each session, volunteers underwent the Attentional Network Task (ANT) and Continuous Compensatory Tracker (CTT) task implemented in the Psychology Experiment Building Language (PEBL) software [46] and administered on a PC monitor (60 cm was the distance from the 27” screen to the eyes, screen resolution 1024 × 768). During the cognitive assessment, participants had to wear the Psychophysiological sEnsoRs Mask FOr Real-Life Cognitive Monitoring (*PERFORM*), a periocular sensorized mask for biosignals acquisition [47]. Thus, the perceived workload was assessed immediately after the completion of the PEBL cognitive tasks for both conditions by using PEBL Nasa Task Load Index (PEBL TLX) scale [46].

### 2.3 Cognitive Assessment

#### 2.3.1 PEBL Attentional Network Task

The Attentional Network Task (ANT) aims at assessing the functioning of alerting, orienting, executive control attentional networks [48,9]. Participants are supposed to determine the direction of the central arrow in a set of five, while ignoring the directions of the surrounding arrows. To indicate the correct direction of the central arrow, participants have to press the corresponding button on the keyboard [49,50]. In the current study, PEBL version of the Attentional Network Test was used and, according to Fan et al. [48], we considered the correct trials (i.e.: not considering the incorrect answers) for the performance indices calculation, represented by: 1) alerting index, 2) orienting index and 3) conflict index.

The efficiency of the alerting network is examined by changes in reaction time (RT) resulting from a warning signal. The alerting index is calculated by subtracting the mean RT of the double-cue condition from the mean RT of the no-cue condition [49].

The efficiency of orienting is examined by changes in the RT that accompany cues indicating the location in which the target will occur. The orienting index is expressed as the difference between the mean RT of the items in a central cue condition (“central cue”) and the average RT of the items in a spatial cue condition (“spatial cue”).

The efficiency of the executive network is examined by requiring the participant to respond by pressing two keys indicating the direction (left or right) of a central arrow surrounded by congruent, incongruent or neutral flankers. The conflict index is calculated by subtracting the mean RT of congruent flanking conditions from the mean RT of incongruent flanking conditions.

#### 2.3.2 PEBL Compensatory Tracker

The Continuous Compensatory Tracking (CCT) is a cognitive test originally developed to assess alertness and vigilance [51,52], and also used to assess sustained attention [53]. Participants have to continuously adjust the position of a pointer to overlap it to a target during eight consecutive trials (from T1 to T8). The pointer is under randomly directed forces that need to be continuously compensated [46]. In the current study, the PEBL version of the CCT task was used to assess vigilance by means of two indices: CCT deviation and CCT speed.

The degree of adaptation was assessed by considering the changes of the deviation and speed from the beginning (T1) to the end (T8) of the task:

- CCT deviation. The median of spatial displacements between the target position and the pointer was calculated for each trial (lower values of median deviation correspond to higher accuracy of task performance) and CCT deviation was estimated as the displacements change from the first to last trial.
- CCT speed. The mean of mouse velocity over the task was calculated for each trial and CCT speed was estimated as the speed change from the first to last trial. Mouse velocity should indicate the degree of the subject’s reactivity toward the task; higher values correspond to a higher degree of reactivity for compensating random motion of the pointer.

#### 2.3.3 *PEBL* Nasa Task Load Index (PEBL TLX)

Nasa Task Load Index [54] is a self-report multidimensional scale aiming at providing an overall perceived workload score, based on a weighted average of six subscales. The subscales are: mental demand, physical demand, temporal demand, own performance, effort and frustration. Subjects have to rate the perceived workload experienced during the previous completion of cognitive tasks by choosing a score ranging from 0 to 100 for the six subscales; higher values indicate greater perceived workload. In the present study, the perceived workload rating was assessed by using the version of NASA TLX on PEBL software (PEBL TLX) [46].

### 2.4 Physiological assessment

During cognitive assessment, participants wore the Psychophysiological sEnsoRs Mask FOr Real-Life Cognitive Monitoring (*PERFORM*), a validated multi-sensorized wearable and non-obtrusive mask [47] able to detect, record and analyze the following physiological signals from a set of dry electrodes placed over the periocular area:

- Facial temperature signals, recorded at 1Hz sampling rate from sensors placed over the left and right zygomatic muscle and the left and right forehead;
- Heart pulse, recorded at 100 Hz sampling rate with a photoplethysmograph sensor placed over glabella (the area between the eyebrows and above the nose);
- Head movements signal recorded at 100 Hz sampling rate from a 3-axial accelerometer placed over the left side of the mask.

For each signal, peripheral measures were extracted to study how performing a cognitive task could change the peripheral outputs. In accordance with the physiological nature of each measurement, we obtained time-series of measures from the beginning to the end of each cognitive task. Thus, as effective parameters associated with the performed task, we considered the changes of each extracted measure from the beginning of the task (average of the measure over the first tenth of its time series) to its end (average of the measure over the last tenth of its time series).

From the facial temperature time-series we considered:

- *MaxT*, defined as the maximum of the four temperature changes calculated between the beginning and the end of the task;
- *zfT*: defined by comparing the aforementioned T changes at the forehead vs those at the cheekbones (*zfT* = *ΔT_Z_ – ΔT_f_* where *ΔT_Z_* is the average changes over the two forehead sensors, and *ΔT_f_* is the average changes over the two cheekbones sensors).

From the heart pulse time-series, we obtained the pulse to pulse time interval series [55], which allowed us to estimate the changes of heart rate HR, defined as the rate change calculated between the beginning and the end of the task.

From the head movements time series, we obtained an integrated measure of head movements from the variance of the combined three axial oscillations calculated within consecutive 1s windows. We estimated the head movement amplitude (HMA) as the changes between the beginning and the end of the task of this measure.

### 2.5 Statistical analysis

Given that this work aims at investigating the physiological correlates of workload of each specific cognitive domain as a function of sleep deprivation, we analyzed the data by following two main steps:

1. identifying sleep deprivation effects on the perceived workload;
2. identifying physiological correlates of intra-subject cognitive performance changes (from baseline to post-sleep deprivation) in the different cognitive tasks after removing the contribution of workload changes.

For step 1), PEBL TLX subscales differences between sleep deprivation and baseline sessions were assessed with a two-tailed Wilcoxon signed-rank test.

For step 2), physiological correlates of intra-subject cognitive performance changes were identified by correlating the changes of cognitive task indices and physiological measures from baseline to post-sleep deprivation. To remove the contribution of workload changes from correlation values between cognitive indices and physiological measures, the partial correlations (partial ranks correlation) were calculated by controlling for those workload subscales that reached the statistical significance at step 1).

The Yekutieli and Benjamini procedure [56] for controlling the false discovery rate (FDR) of the family of hypothesis tests concerning all physiological features correlated with each cognitive task index was applied. The false discovery rate was set equal to 0.05 and adjusted p values were calculated.

## 3. Results

### 3.1 Perceived workload changes from baseline to sleep deprivation

The perceived workload measured after tasks was different between conditions for the mental and physical demand subscales: mental and physical demands increased in post-sleep deprivation condition (p=0.004 and p=0.007, respectively as well as p_adj_=0.025 and p_adj_=0.025, respectively). Table 1 provides statistics of each PEBL TLX subscales.

**Table 1:** PEBL Task Load Index

| | condition | $m \pm s.e.$ | $\Delta (m \pm s.e.)$ | Signed Rank | $p$-value |
| --- | --- | --- | --- | --- | --- |
| Mental Demand | B | $27 \pm 4.82$ | $-11.66 \pm 5$ | 24 | .004* |
| | D | $39 \pm 5.42$ | | | |
| Physical Demand | B | $18 \pm 2.71$ | $-21 \pm 7$ | 9 | .007** |
| | D | $39 \pm 6.62$ | | | |
| Temporal Demand | B | $43 \pm 6$ | $-5.33 \pm 8.41$ | 46.5 | ns |
| | D | $48 \pm 7$ | | | |
| Frustration | B | $39 \pm 4.29$ | $-3.66 \pm 5.46$ | 35.5 | ns |
| | D | $42 \pm 4.36$ | | | |
| Effort | B | $33 \pm 5.36$ | $-14.66 \pm 8$ | 27.5 | ns |
| | D | $48 \pm 7$ | | | |
| Performance | B | $24 \pm 6$ | $-5.66 \pm 10$ | 36 | ns |
| | D | $30 \pm 7$ | | | |
| Overall Workload | B | $31 \pm 2$ | $-10 \pm 5$ | 95 | ns |
| | D | $41 \pm 4.35$ | | | |

B=baseline condition, D=sleep deprived condition; \* Wilcoxon signed rank test with $p < 0.05$ ; \*\* Wilcoxon signed rank test with $p < 0.01$ ;

### 3.2 Physiological correlates of attention systems functioning

None of the indices measuring the cognitive performance during the tasks were significantly changed after sleep deprivation. However, significant associations were found between cognitive indices and physiological measures after removing the putative linking effect related to the changes of perceived workload.

As regards ANT, the subjects who displayed an increased alerting score in post-sleep deprivation condition showed a decreased HMA: in fact, a negative correlation was found between ANT alerting index and the HMA measure after correction for PEBL TLX mental and physical demands (r_p_=-0.73, p=0.0025, p_adj_=0.01).

As regards CCT, the subjects who displayed an increased CCT speed in post-sleep deprivation condition showed increased variation of temperature. After correction for mental and physical demands, a positive correlation was found between CCT speed and *zfT*(r_p_=0.74, p=0.0015, p_adj_=0.006) and between CCT speed and maxT (r_p_=0.66, p=0.0067, p_adj_=0.013).

## 4. Discussion

Our study aimed at identifying peripheral physiological correlates of attention components in healthy subjects. A within-subject study using sleep deprivation as a stressor [13] was used to induce changes in individual cognitive performances. Each participant underwent two experimental sessions, both conducted at 6 pm, and one of which took place after one night of sleep deprivation. For each session, physiological signals were recorded using the *PERFORM* [47] during the completion of two cognitive tasks assessing attentional networks efficiency. After cognitive tests, an evaluation of the perceived workload was performed. The evaluation of perceived workload allowed us to get an estimate of its contribution to cognitive functioning changes from the baseline to sleep deprivation condition and to remove its unspecific contribution to the physiological/cognitive relationships.

Herein, physiological signals were derived from peri-ocular sites by using the *PERFORM* [47], a system conceived to be used in VR headset.

### 4.1 Sleep deprivation increases mental and physical demands

Our results highlighted an increased mental and physical demand in post-sleep deprivation condition as compared to baseline. Several studies reported that the progress of prolonged wakefulness leads to a gradual increase in the homeostatic biological drive [15,57,58] that consequently augments the state of sleepiness. In this context, perceived workload increase after a period of prolonged wakefulness is reported [59,60] and explained as a counterbalancing response to a fatigued state [16]. Liu and colleagues [61] reported a general increase in all workload subscales after 32 hours of sleep deprivation in a sample of expert airplane pilots after simulated flight tasks. Tomasko and colleagues [59] assessed workload in sleep-deprived medical students after laparoscopic-simulated tasks while Fairclough and colleagues [62] did the same in sleep-deprived subjects after primary driving tasks and both highlighted increased scores for all NASA-TLX subscales, except for the mental demand one.

Our findings partially agree with the previous studies, since perceived workload increase was limited to mental and physical demand subscales. It is worth noting that previous studies enrolled selected group of participants for their investigations, thus administering tasks intimately linked to the subjects’ activities of daily living (i.e. surgical task for surgeons, simulator flight task for expert flyers), whereas, in the present research, tasks were not part of everyday life and they could represent a novel experience. The novelty of the tasks, the effort to comprehend instructions, the lack of similarity with activities of daily living could represent additional mental stressors that are exacerbated by sleep deprivation. However, it is worth noting that the scores of cognitive tasks did not show significant differences between conditions (baseline and post-sleep deprivation): in this context, it is possible to suggest that the higher the perceived workload, the higher the effort to counteract fatigue to guarantee an adequate cognitive performance.

### 4.2 The physiological profile differs between cognitive functions

Our results highlighted the associations between changes from baseline to post-sleep deprivation condition of physiological measures and cognitive indices. These associations were putatively sustained by different factors: the general increase of workload and the modulation of the cognitive functions specifically involved in each task. Most of the associations identified between physiological measures and cognitive indices held true also after mental and physical demands correction, suggesting a strong contribution of specific cognitive functions on physiological reactions.

As regards the attentional domain, a higher reactivity of the alerting system in post-sleep deprivation condition was associated with a decreased amplitude of head movements. Accordingly, a study of head movements during surgical tasks has shown that fewer head movement amplitude during a laparoscopic simulation was associated with better learning [63]. Our results sustain the hypothesis that fewer head movements are related to more precise and more focused performances during tasks.

Concerning alertness and vigilance during visuo-motor tracking task, a higher reactivity (CCT speed) after sleep deprivation was shown in the participants who displayed greater temperature changes from the beginning to the end of the task between cheekbones and forehead sensors (*zfT*) and between all facial sensors. Classically, temperature fluctuations during tasks are associated with task performance. It has been reported that brain metabolism increase during cognitive tasks implies more heat production [23]. As some blood vessels connect facial tissues with the brain [64,65] metabolic brain changes cause variations in the peripheral skin temperature that we were able to detect.

## 5. Conclusions

This integrated evaluation of cognitive functioning using subjective, behavioral, and physiological measures allowed us to gain a better comprehension of attentional systems and their relative physiological changes after sleep deprivation. An augmented perceived physical and mental demand was shown in post-sleep deprivation condition. Head movements and temperature variations were revealed to be sensitive to changes occurring during specific cognitive performance in sleep deprivation as compared to baseline.

**Figure 2:**
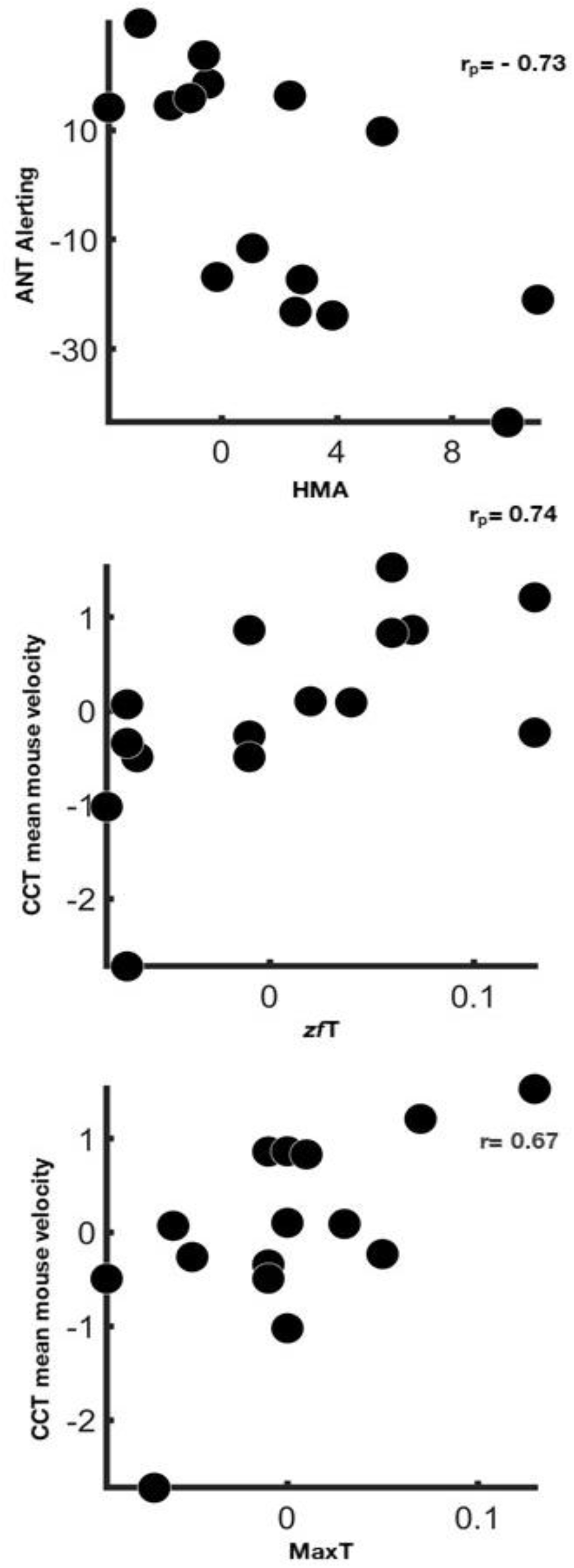
Physiological correlates of attention systems functioning. Scatter plots show associations between physiological measures and cognitive indices changes from baseline to post-sleep deprivation condition after the correction for TLX mental and physical demand (r_p_, partial rank correlation coefficient - False Discovery Rate<0.05).

## Contributions

Valentina Cesari: Methodology, Investigation, Writing - Original Draft, Visualization, Writing - Reviewing and Editing;

Elena Marinari: Investigation, Writing - Original Draft, Visualization, Resources;

Marco Laurino: Software, Writing - Original Draft, Writing-Reviewing and Editing;

Angelo Gemignani: Conceptualization, Supervision, Original Draft, Writing-Reviewing and Editing;

Danilo Menicucci: Conceptualization, Supervision, Formal Analysis, Methodology, Visualization, Original Draft, Writing-Reviewing and Editing.

Valentina Cesari and Elena Marinari contributed equally to this manuscript

## Ethics and Disclosure Statement

All participants signed informed consent and the study protocol was approved by the Committee on Bioethics of the University of Pisa (Review No. 16/2019 Meeting held on June 28, 2019).

Financial Disclosure Statement: The research was partially funded by the Project “Brain Machine Interface in space manned missions: amplifying Focused attention for error Counter-balancing” (BMI-FOCUS, Tuscany Region POR CREO 2014/2020).

Nonfinancial Disclosure Statement: The authors report no nonfinancial conflicts.

## Acknowledgements

We thank Davide Cini and Andrea Berton (both at the Institute of Clinical Physiology – CNR Pisa) for the technical assistance. We also thank Eleonora Malloggi for the revision of English language.

